# Real-Time Search Assisted Acquisition on a Tribrid Mass Spectrometer Improves Coverage in Multiplexed Single-Cell Proteomics

**DOI:** 10.1101/2021.08.16.456445

**Authors:** Benjamin Furtwängler, Nil Üresin, Khatereh Motamedchaboki, Romain Huguet, Daniel Lopez-Ferrer, Vlad Zabrouskov, Bo T. Porse, Erwin M. Schoof

**Affiliations:** The Finsen Laboratory, Rigshospitalet, Faculty of Health Sciences, University of Copenhagen, Copenhagen, Denmark; Biotech Research and Innovation Centre (BRIC), University of Copenhagen, Copenhagen, Denmark; Novo Nordisk Foundation Center for Stem Cell Biology, DanStem, Faculty of Health Sciences, University of Copenhagen, Copenhagen, Denmark; Department of Biotechnology and Biomedicine, Technical University of Denmark, Lyngby, Denmark; ThermoFisher Scientific, San Jose, CA, USA

**Author notes:** Corresponding Authors: Benjamin Furtwängler, Erwin M. Schoof.

**Keywords:** single-cell proteomics, isobaric tag quantification, multiplexing, SPS-MS3, TMT, real-time-search

## Abstract

In the young field of single-cell proteomics (scMS), there is a great need for improved global proteome characterization, both in terms of proteins quantified per cell and quantitative performance thereof. The recently introduced real-time search (RTS) on the Orbitrap Eclipse Tribrid mass spectrometer in combination with SPS-MS3 acquisition has been shown to be beneficial for the measurement of samples that are multiplexed using isobaric tags. Multiplexed single-cell proteomics requires high ion injection times and high-resolution spectra to quantify the single-cell signal, however the carrier channel facilitates peptide identification and thus offers the opportunity for fast on-the-fly precursor filtering before committing to the time intensive quantification scan. Here, we compared classical MS2 acquisition against RTS-SPS-MS3, both using the Orbitrap Eclipse Tribrid MS with the FAIMS Pro ion mobility interface and present a new acquisition strategy termed RETICLE (**R**TS **E**nhanced Quan**t** of S**i**ngle **C**el**l** Sp**e**ctra) that makes use of fast real-time searched linear ion trap scans to preselect MS1 peptide precursors for quantitative MS2 Orbitrap acquisition. We show that classical MS2 acquisition is outperformed by both RTS-SPS-MS3 through increased quantitative accuracy at similar proteome coverage, and RETICLE through higher proteome coverage, with the latter enabling the quantification of over 1000 proteins per cell at a MS2 injection time of 750ms using a 2h gradient.

## INTRODUCTION

Recent developments in liquid chromatography coupled mass spectrometry (LC-MS) based proteomics revealed the high potential for its application on single cells^1–9^. A major breakthrough was the introduction of single-cell proteomics using mass spectrometry (scMS) via the SCoPE^1,2,10^ method, where isobaric labeling is used to multiplex single-cells which are then measured in a single LC-MS run. Importantly, the addition of a carrier channel (200-cell equivalent) provides significantly more peptides copies in addition to the single-cell channels and thus facilitates precursor ion identification in the fragmentation scan (MS2). Hence, the increased throughput permitted through isobaric multiplexing, which currently supports up to 18 channels with TMTpro^11,12^, and the signal boosting effect of the carrier channel, provide considerable advantages over label-free scMS. Nonetheless, multiplexed scMS suffers from its own limitations. For one, the carrier channel decreases the quantitative performance of the adjacent channels due to signal spillover from isotopic impurities of the isobaric labels and furthermore increases the overall noise level relative to the very low abundant single-cell channels^5,9,13,14^. Moreover, long injection times are required to collect enough ions from the single-cell channels for robust ion statistics and accurate reporter ion quantification, resulting in slow scanning speed and consequently lower proteome coverage. Moreover, isobaric reporter ion quantification suffers from the well-studied problem of ratio-compression due to interference of co-isolated precursor ions^15,16^. This interference can be mitigated by applying gas-phase fractionation via the FAIMS interface^17,18^, which reduces the complexity of the precursor ion stream entering the MS and removes +1 ion species (i.e. non-peptide contaminants). Even greater reduction of interference is achieved with the gas-phase purification of the MS2 fragment ions and subsequent secondary fragmentation to release the isobaric tag for quantification in a MS3 scan^16^. This feature is unique to Tribrid MS instruments as synchronous precursor selection (SPS) is performed in the linear ion trap (LIT). Since the high resolution scan in the Orbitrap (OT) is only needed to measure accurate precursor masses in the MS1 scan and to resolve the reporter ions for quantification in the MS3 scan, peptide identification via MS2 can be performed in the more sensitive LIT^19^. This results in fast and sensitive MS2 acquisition, as the Tribrid design enables parallelization of OT and LIT scans. Additionally, with the introduction of real-time search (RTS)^20,21^ on the Orbitrap Eclipse Tribrid MS^22^, only MS1 precursors that were identified as peptides of interest are subjected to the time consuming MS3 quantification, resulting in higher proteome coverage. Furthermore, since the MS1 precursor is identified, only the peptide fragments belonging to that peptide are subjected to SPS-MS3, reducing the co-isolation to a minimum and thus maximizing accuracy. Previously, SPS-MS3 without RTS was applied to scMS^5^, resulting in much lower proteome coverage; a limitation that RTS could overcome.

We recently presented our scMS workflow^7^ and benchmarked the quantitative performance using the OCI-AML8227 cell-culture model^23^. This model maintains the hierarchical nature of Acute Myeloid Leukemia (AML) where a small population of self-renewing leukemic stem cells (LSC) differentiate to progenitors (PROG) and finally to terminally differentiated blasts (BLAST). Thus, this model system provides three distinct cell differentiation stages, all contained in one cell culture, with differences on proteome level detectable by scMS. Furthermore, we investigated the protein fold-changes between these differentiation stages via bulk proteomics to a depth of nearly 7000 proteins, providing us with a reference set to benchmark scMS data in terms of quantitative accuracy.

In this work, we compared the performance of multiplexed scMS using MS2 acquisition against RTS-SPS-MS3, both using the Orbitrap Eclipse Tribrid MS with the FAIMS Pro ion mobility interface. Furthermore, we present a new acquisition strategy termed RETICLE (**R**TS **E**nhanced Quan**t** of S**i**ngle **C**el**l** Sp**e**ctra) that makes use of fast real-time searched linear ion trap scans to preselect MS1 peptide precursors for quantitative MS2 Orbitrap acquisition. We show that classical MS2 acquisition is outperformed both by RTS-SPS-MS3 through increased quantitative accuracy at similar proteome coverage, and by RETICLE through higher proteome coverage, with the latter enabling the quantification of over 1000 proteins per cell at a MS2 injection time of 750ms using a 2h gradient.

## EXPERIMENTAL PROCEDURES

### Cell Culture and FACS Sorting

OCI-AML8227 cells were grown in StemSpan SFEM II media, supplemented with growth factors (Miltenyi Biotec, IL-3, IL-6 and G-CSF (10 ng/mL), h-SCF and FLt3-L (50 ng/mL), and TPO (25 ng/mL) to support the hierarchical nature of the leukemia hierarchy captured within the cell culture system. On day 6, cells were harvested (8e6 cells total), washed, counted, and resuspended in fresh StemSpan SFEM II media on ice at a cell density of 5e6 cells/ml. Staining was done for 30 mins on ice, using a CD34 antibody (CD34-APC-Cy7, Biolegend, clone 581) at 1:100 (vol/vol), a CD38 antibody (CD38-PE, BD, clone HB7) at 1:50 (vol/vol). Cells were washed with extra StemSpan SFEM II media, and subsequently underwent three washes with ice cold PBS to remove any remaining growth factors or other contaminants from the growth media. Cells were resuspended for FACS sorting in fresh, ice cold PBS at 2e6 cells/ml and stained with 7-AAD viability dye (1 ug/mL, Invitrogen). Cell sorting was done on a FACSAria III instrument, controlled by the DIVA software package (v.8.0.2) and operating with a 100 μm nozzle. Cells from 3 different gates (CD34+CD38-, CD34+CD38+, CD34-) (**Supplementary Fig. 1**) were sorted at single-cell resolution into a 384-well Eppendorf LoBind PCR plate (Eppendorf AG) containing 1 μl of lysis buffer (50 mM Triethylammonium bicarbonate (TEAB) pH 8.5, 20% 2,2,2-Trifluoroethanol (TFE)). Directly after sorting, plates were briefly spun, snap-frozen on dry ice, and then boiled at 95 °C in a PCR machine (Applied Biosystems Veriti 384-well) for 5 mins. Plates were again snap-frozen on dry ice and stored at -80 °C until further sample preparation. The same procedure was followed for the carrier plate, but instead of sorting single cells, 500 cells were sorted in four-way purity mode into each well without immunofluorescent preselection.

### Sample Preparation of Diluted Standard

Peptide concentration of the TMTpro labeled OCI-AML8227 sample that was used to measure the MS3 reference library as previously described^7^ was determined via Nanodrop and the sample was subsequently diluted to contain 250 pg of peptide in each of the 9 channels per injection. A bulk-sorted OCI-AML8227 carrier was added as a 200-cell equivalent per injection. For the comparison of the close-out options, samples with a 100-cell equivalent carrier were used.

### Sample Preparation of Single-Cell Samples

After thawing, protein lysates from the single cells were digested with 2 ng of Trypsin (Sigma cat. nr. T6567), dissolved in 1 μl of 100 mM TEAB pH 8.5 containing Benzonase (Sigma cat. nr. E1014) diluted 1:5000 (vol/vol) to digest any DNA that would interfere with downstream processing. For the carrier plates, the amount of trypsin was increased to 10 ng in order to digest the protein content of each well containing 500 cells. Plates were kept at 37 °C overnight to complete the protein digestion. All dispensing steps in this protocol were done using the Dispendix I-DOT One instrument. After digestion, peptides were labeled with TMTPro reagents. 6 mM in 1 μl acetonitrile (ACN) of each label was added to the single-cell wells, while the 500-cell carrier plate wells were labeled with 13 mM of TMTPro-126 reagent in each well. Subsequently, plates were kept at RT for 1 h. The labeling reaction was quenched with 1 μl of 1.25% or 2.5% Hydroxylamine for the single-cell and carrier plate respectively for 15 min. Subsequently, the carrier plate was pooled and desalted using a SOLAμ HRP 96-well plate. Eluted and desalted peptides were concentrated to dryness in an Eppendorf Speedvac, after which they were resuspended in A* (2% ACN, 0.1% TFA. The single-cell samples were then mixed from 14 single cells plus the equivalent of 200 carrier channel cells. To ensure a balanced design, each sample contained 4 cells of CD34- and 5 cells of CD34+CD38- and CD34+CD38+ respectively. This pooling was performed using the Opentrons OT-2 liquid handler. The resulting peptide mix was concentrated in an Eppendorf Speedvac, and re-constituted in A* for individual Mass Spectrometry analysis.

### Mass Spectrometry Data Acquisition of Diluted Standard

Peptides were loaded onto a µPAC trapping column (PharmaFluidics) at 5 ul/min, connected in a forward-flush configuration to a 50 cm µPAC analytical column (PharmaFluidics), with 100% Buffer A (0.1% Formic acid in water) using the UltiMate 3000 RSLCnano System (ThermoFisher), and the column oven operating at 35 °C. Peptides were eluted over a 90 min gradient, using 80% Acetonitrile, 0.1% Formic acid (Buffer B) going from 8% to 18% over 8 min at a flowrate of 500 nl/min, to 19% over 1 min with a simultaneous reduction of flowrate down to 150 nl/min, which was kept at this rate from there on. Then to 28% over 31 min, to 38% over 20 min, to 60% over 6 min, to 95% over 1 min, and kept at 95% for 23 min, after which all peptides were eluted. Spectra were acquired with an Orbitrap Eclipse Tribrid Mass Spectrometer with FAIMS Pro Interface (ThermoFisher Scientific) running Tune 3.4 and Xcalibur 4.3. For all acquisition methods, FAIMS switched between CVs of -50 V and -70 V with cycle times of 3s and 1.5s, respectively. MS1 spectra were acquired at 120,000 resolution with a scan range from 375 to 1500 m/z, normalized AGC target of 300% and maximum injection time of 50ms. Precursors were filtered using monoisotopic peak determination set to peptide, charge state 2-6, dynamic exclusion of 120s with ±10 ppm tolerance excluding isotopes and different charge states, and a precursor fit of 70% in a windows of 0.7 m/z for MS2 and RETICLE and 60% in a 0.8 m/z window for RTS-MS3. Additionally, RTS-MS3 employed an intensity threshold of 5e3. Precursors selected for MS2 analysis were isolated in the quadrupole with a 0.7 m/z window for MS2 and RETICLE and 0.8 m/z for RTS-MS3. In the MS2 method, ions were collected for a maximum injection time of 500ms or 750ms and normalized AGC target of 500%, fragmented with 32 normalized HCD collision energy and MS2 spectra were acquired in the Orbitrap at 120,000 resolution with first mass set to 120 m/z. In the RTS-MS3 and RETICLE methods, ions were collected for a maximum injection time set to auto for RTS-MS3 and 23ms for RETICLE and normalized AGC target of 300% for both methods. Fragmentation was performed with 30 normalized CID collision energy and MS2 spectra were acquired in the LIT at scan rate rapid for RTS-MS3 and turbo for RETICLE. For the RETICLE method, LIT MS2 spectra were subjected to RTS using the homo sapiens database obtained from Uniprot (Swiss-Prot with isoforms) and Trypsin set as enzyme. Static modifications were TMTpro16plex on Lysine (K) and N-Terminus, and Carbamidomethyl on Cysteine (C). Oxidation on Methionine (M) was set as variable modification. Maximum missed cleavages was set to 0 and maximum variable mods to 1. FDR filtering was enabled, maximum search time was set to 40ms, and the scoring threshold was set to 1.4 XCorr, 0.1 dCn and 10 ppm precursor tolerance. Use as trigger only was checked and close-out was enabled with maximum number of peptides per protein set to 4. For the comparison of the close-out options, this number was either set to 4, 10 or close-out was disabled. Precursors identified via RTS were isolated in the quadrupole with a 0.7 m/z window, ions were collected for a maximum injection time of 500ms or 750ms and normalized AGC target of 500%, fragmented with 32 normalized HCD collision energy and MS2 spectra were acquired in the Orbitrap at 120,000 resolution with first mass set to 120 m/z. For the RTS-MS3 method, LIT MS2 spectra were subjected to RTS using the same settings as in RETICLE with differences being that maximum missed cleavages was set to 1, maximum variable mods to 2, maximum search time to 35ms and use as trigger only was deactivated, but TMT SPS MS3 Mode was activated. Subsequently, spectra were filtered with a precursor selection range filter of 400-1600 m/z, isobaric tag loss exclusion of TMTpro and precursor mass exclusion set to 5 m/z low and 3 m/z high. Precursors identified via RTS were isolated for a MS3 scan using the quadrupole with a 2 m/z window and ions were collected for a maximum injection time of 500ms or 750ms and normalized AGC target of 500%. SPS was activated and number of SPS precursors was set to 10. Isolated fragments were fragmented again with 45 normalized HCD collision energy and MS3 spectra were acquired in the Orbitrap at 120,000 resolution with a scan range of 100-500 m/z. Method trees are shown in **Supplementary Fig. 2**.

### Mass Spectrometry Data Acquisition of scMS Samples

Real scMS samples were measured as described for the diluted standard with the following changes. Peptides were eluted over a 116 min gradient, using 80% Acetonitrile, 0.1% Formic acid (Buffer B) going from 8% to 15% over 8 min at a flowrate of 500 nl/min, to 15.6% over 1 min with a simultaneous reduction of flowrate down to 150 nl/min, which was kept at this rate from there on. Then to 19% over 12 min, to 27% over 45 min, to 38% over 26 min, to 95% over 1 min, and kept at 95% for 23 min, after which all peptides were eluted. Cycle times for FAIMS switching were set to of 2.5s and 1.2s for -50 and -70 CV, respectively. For RTS, maximum number of peptides was set to 5 and all methods used maximum number of missed cleavages of 1 and maximum variable mods of 2. Carbamidomethyl on Cysteine (C) was not set, as these samples were not treated with TCEP/CAA.

### Mass Spectrometry Raw Data Analysis

Raw files were analyzed with Proteome Discoverer 2.4 (ThermoFisher Scientific) with the built-in TMTPro Reporter ion quantification workflows using the standard settings if not further specified. Spectra were searched using the Sequest search engine using the homo sapiens database obtained from Uniprot (Swiss-Prot with isoforms). Static modifications were TMTpro16plex on Lysine (K) and N-Terminus, and for the diluted standard Carbamidomethyl on Cysteine (C) was set. Dynamic modifications were set as Oxidation (M), and Acetyl and Met-loss on protein N-termini. Fixed modifications were set to TMTPro on both peptide N-termini and K residues. Results were re-scored with Percolator and filtered to 1% FDR. For the MS2 and RETICLE methods, reporter ion quantification was performed on FTMS MS2 spectra and the same spectra were also sent to Sequest for identification, where they were searched with fragment mass tolerance of 0.02 Da. For the RTS-MS3 method, reporter ion quantification was performed on FTMS MS3 spectra and LIT MS2 spectra were searched with fragment mass tolerance of 0.6 Da. For the reporter ion quantification in all methods, normalization mode and scaling mode were set to None and average reporter s/n threshold was set to 0. Isotopic error correction was applied.

### Data Analysis of Diluted Standard

Proteome Discoverer 2.4 result tables were loaded into python 3.7.10. Potential contaminant proteins were removed. The first three “single-cell” channels (127N, 128N, 129N) were excluded from the analysis as they might be contaminated from the isotopic impurities of the carrier channel (126). Protein S/N values below 1.1 were set to missing values. Technical triplicates of each of the six methods were normalized by equalizing the median s/n of the “single-cell” channels for each protein across replicates (one correction factor per protein). Differential protein expression (DE) analysis was performed on normalized and log2 transformed data with a two-sided Welch’s t-test, excluding missing values. P-values were corrected via the Benjamini–Hochberg procedure and a cutoff of 5% FDR was applied. Proteins were testable if each group contained at least three values. Proteome Discoverer calculated Log2FC and adj.pval for the bulk-measured reference. True positive (TP) DE proteins were defined to have less than 5% adj.pval in both scMS and the bulk-measured reference and the same log2FC directionality. False positive (FP) DE protein were defined to have less than 5% adj.pval in scMS but different directionality or >5% adj.pval in the reference. True negative (TN) were defined to have >5% adj.pval in both scMS and reference. False negative (FN) were defined to have >5% adj.pval in scMS and <5% adj.pval in the reference. FDR of DE detection was defined as FP/(TP+FP), sensitivity as TP/(TP+FN) and specificity as FP/(TN+FP).

### Data Analysis of scMS Samples

scMS data was analyzed as previously described^7^. Briefly, FACS.fcs files were processed in FlowJo 10.7.1 with the IndexSort 2.7 plugin to apply bi-exponential transform. FACS data and sort- and label layouts were used to create the metadata for each cell. Subsequent analysis was performed with python 3.7.10 and SCeptre version 1.0, a python package that extends the functionalities of Scanpy^24^. First, SCeptre normalization and cell filtering were applied. Subsequently, proteins were filtered to be quantified in at least 10 cells and data was log2 transformed. This data was used for figures 3A-C & E. For figures 3D & F, data was imputed using the k-nearest neighbor algorithm and protein expression was scaled. The silhouette coefficients were calculated using the ‘silhouette_samples’ function from Scikit-learn, providing the population labels and the first 10 principal components as matrix.

### Experimental Design and Statistical Rationale

The diluted standard was measured as technical triplicate to include technical variance between injections and stochastic peak sampling from data dependent acquisition. For the real scMS samples, over 110 single-cells were measured for each method representing biological replicates and thus including biological variance. For DE analysis, at least three values were required in each of the two compared groups to calculate log2FC and p-value and the latter were corrected via the Benjamini–Hochberg procedure.

## RESULTS

### Design of Acquisition Methods

We created three different acquisition methods (MS2, RTS-MS3, RETICLE) for the Orbitrap Eclipse Tribrid Mass Spectrometer via the Xcalibur Instrument Setup (**Supplementary Fig. 2**). In the MS2 method, precursors for the quantitative MS2 scan are directly selected from the MS1 scan. In the RTS-MS3 method, precursors detected in the MS1 scan are subjected to an MS2 scan in the LIT, which is subsequently searched against a user-defined database. If the precursor is identified as a peptide of interest, it is subjected to a quantitative SPS-MS3 scan. The same procedure is applied in RETICLE, with the difference that precursors identified as a peptide of interest are subjected to a quantitative MS2 scan. Thus, MS2 and RETICLE acquisition yields high-resolution HCD-OT-MS2 scans that are used for peptide identification and quantification in the subsequent data analysis, whereas RTS-MS3 yields low-resolution CID-LIT-MS2 scans for peptide identification and linked high-resolution HCD-OT-MS3 scans that can only be used for quantification. To investigate how the ion injection time (IT) of the quantitative scan influences proteome coverage and quantitative performance, we tested each acquisition method with IT of 500ms and 750ms resulting in six methods (MS2 500ms, MS2 750ms, RTS-MS3 500ms, RTS-MS3 750ms, RETICLE 500ms, RETICLE 750ms).

### Comparison of Acquisition Methods Using a Diluted Standard

To enable a reproducible comparison between MS acquisition methods we created a multiplexed sample that can be injected as technical replicate but also preserves the biologically relevant protein fold-changes between LSC, PROG, and BLAST. For this, the three differentiation stages were FACS sorted and equal numbers of cells were labeled with three TMTpro channels per differentiation stage. Subsequently, the sample was diluted to contain 250 picogram of peptides per “single-cell” channel per injection and a carrier channel was added as a 200-cell equivalent (**Fig 1A**). This sample was measured in triplicates for each of the six methods. The analysis of the raw files showed that with RTS-MS3 and RETICLE, less quantification scans were recorded (**Fig 1B**). Nonetheless, these RTS-triggered scans had considerably higher identification rates, resulting in more quantified proteins in the “single-cell” channels when comparing the same ITs. The higher identification rate in RETICLE compared to RTS-MS3 showed the advantage of using the high-resolution long-injected HCD-OT-MS2 scans for peptide identification and simultaneous quantification, as this resulted in the highest protein coverage. Nonetheless, RTS-MS3, with its reliance on CID-LIT spectra for peptide identification, performed similar to the MS2 method in terms of protein coverage. Another benefit of RTS is the possibility to limit the number of peptides per protein on the fly using the close-out filter, which improved the distribution of quantified peptides across proteins by reducing the fraction of proteins with only one peptide or with disproportionally many peptides (e.g. over 7 peptides) (**Fig. 1C**).

**Figure 1.**
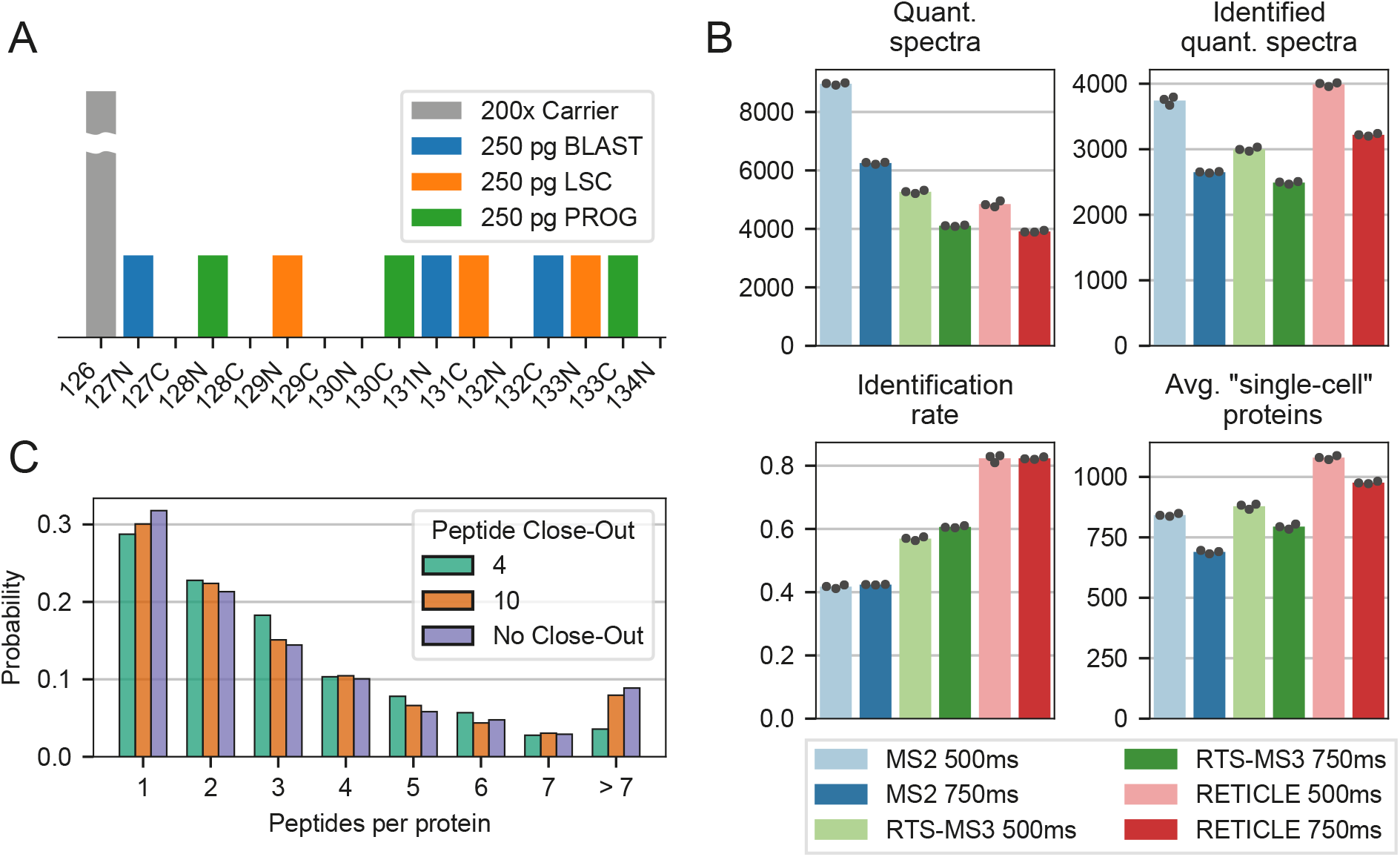
Comparison of scMS performance between the acquisition methods. A) Design of the multiplexed diluted standard. B) Number of acquired quantification spectra differs between the methods, as does the identification rate thereof, influencing the average number of proteins quantified in the “single-cell” channels. C) Close-out option implemented in RTS influences the distribution of peptides across proteins.

The effect of the acquisition method on the resulting protein expression matrix became apparent when ranking the quantified proteins by their S/N (summed peptide S/N), a proxy for protein abundance (**Fig. 2A, Supplementary Fig. 3**). Whereas higher abundant proteins tended to be quantified across all methods, the overlap decreased for lower abundant proteins. For the MS2 method, increased ITs resulted in decreased proteome coverage of lower abundant proteins, however improved S/N for higher abundant proteins and thus exemplified the tradeoff of distributing the acquisition time across more peptides to increase proteome depth while decreasing the quantitative performance. For the RETICLE method, proteome coverage was greatly increased compared to MS2, indicating that RTS in combination with the close-out filter effectively distributed the acquisition time across more proteins. In addition, in both RTS methods, higher IT decreased proteome coverage only slightly while improving S/N of all proteins, indicating that these RTS methods enable the use of very high ITs. To evaluate the quantitative performance of the different methods, we performed differential protein expression (DE) analysis between BLAST and LSC using the bulk-measured reference dataset for validation (**Fig. 2B**). The results showed that RETICLE 750ms and RTS-MS3 750ms provided the highest number of true positive DE proteins. Here, RTS-MS3 750ms provided less proteins to be tested, however outperformed RETICLE 750ms in FDR, sensitivity and specificity. Moreover, although the higher ITs decreased the testable proteins in the RTS methods, true positive DE proteins increased due to improved FDR, sensitivity and specificity. Analyzing the cumulative distribution of detected DE proteins over the abundance range revealed that highly abundant DE proteins were detected at a similar rate for all methods and differences only arose at decreasing abundance (**Fig. 2C**). Here, MS2 750ms likely underperformed since these lower abundant proteins were not sampled, whereas RETICLE 500ms underperformed across a wide abundance range, indicating inferior protein quantification due to the distribution of acquisition time over more proteins. RTS-MS3 750ms outperformed RETICLE 750ms up to the lower abundant proteins, likely due to the improved quantitative accuracy, until the advantages of the increased proteome depth of RETICLE 750ms evened out the results. To investigate the extend of ratio compression, we calculated ratios between fold-changes from scMS and the bulk-measured reference dataset (**Fig. 2D**). The results showed that log2 fold-changes tended to be systematically lower in MS2 methods with median around 20% compared to RTS-MS3 with a median around 5%. Furthermore, we found that quantitative accuracy decreased with lower protein S/N (**Fig. 2E**), however RTS-MS3 outperformed the MS2 methods across the whole intensity range. Taken together, these results suggest that RTS-MS3 750ms outperforms the classical MS2 method by providing more accurate protein quantification at similar proteome coverage and RETICLE 750ms outperforms classical MS2 through higher proteome coverage. In addition, both methods benefit from the improved distribution of acquisition time across proteins enabled by RTS.

**Figure 2.**
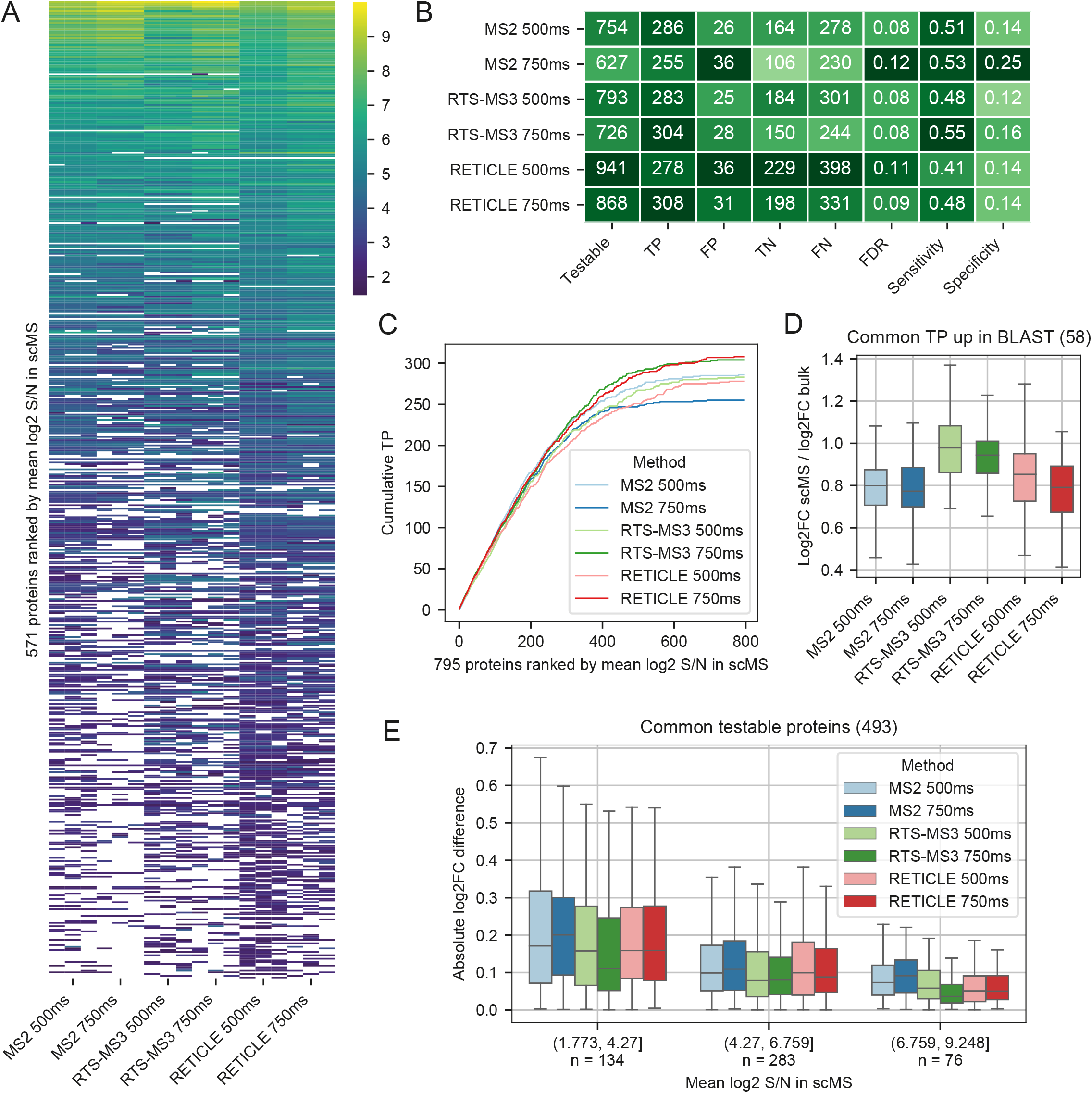
Comparison of acquisition methods using a diluted standard. A) Protein expression matrix of each method measured in triplicates. Proteins in the rows were sorted by the mean log2 S/N across all methods and each individual row shows the same protein with its mean log2 S/N across all “single-cells” in each LC-MS run. Heatmap was downsampled by removing every second row to visualize individual missing values. Full heatmap shown in Supplementary Fig. 3. B) Differential protein expression analysis between BLAST and LSC of each method using the bulk-measured reference dataset for validation. TP=true positive, FP=false positive, TN=true negative, FN=false negative (see Methods). C) Cumulative distribution of true positive DE proteins ordered by mean log2 S/N across all methods. Proteins on the x-axis are the same for each method and represent the union of all testable proteins across all methods that are DE in the bulk-measured reference. D) Ratio compression measured by dividing log2FC in scMS by log2FC in the bulk-measured reference. Common true positive proteins that were upregulated in BLASTS were used. Outlier not shown. E) Absolute log2FC difference between scMS and the bulk-measured reference. The intersection of testable proteins across all methods was used and binned by mean log2 S/N across all methods. Outlier not shown.

### Comparison of Acquisition Methods Using scMS Samples

To investigate the performance of the different acquisition methods with real scMS samples, we measured over 110 single cells for each of the six methods. For this, we FACS sorted single cells from three different populations (CD34+CD38-, CD34+CD38+, CD34-) (**Fig. 3A**) from the OCI-AML8227 cell-culture system into 384-well plates and prepared the cells as previously described^7^. After data acquisition, each dataset was processed with our scMS data analysis pipeline SCeptre^7^, which includes data normalization, cell filtering, imputation and embedding. The results showed that with RETICLE, considerably more proteins were identified across the whole dataset, whereas RTS-MS3 identified similar numbers of proteins as MS2 (**Table 1**). Furthermore, RETICLE 750ms measured the highest number of high-coverage proteins, even surpassing MS2 500ms. When comparing the number of proteins measured in each cell (**Fig. 3B**), it became apparent that RETICLE outperformed MS2 by 35% (MS2 750ms: 711, RETICLE 750ms: 962; median proteins per cell) and RTS-MS3 performed similarly to MS2. Furthermore, the number of proteins per cell in the RTS methods did not change much between ITs, indicating that even higher ITs could be feasible. When comparing measured protein S/N in each dataset (**Fig. 3C**), it became apparent that RETICLE 750ms had the overall highest summed S/N, the most beneficial distribution of S/N across proteins and that the additional proteins had similar S/N values as low ranked proteins in the other methods. DE analysis revealed that DE proteins were mainly detected at higher S/N and differences only arose at decreasing abundance with similar observations as with the diluted standard (**Fig. 3C**). To further evaluate the quantitative performance of proteins measured with low S/N values, we measured how well the populations were separated based on scMS data using principle component analysis (PCA) (**Fig. 3D**). The results showed that separation was equally possible with proteins with lower S/N as with higher S/N. Thus, although detection of DE proteins was more challenging at low S/N, these proteins provided enough signal to separate different cell populations, indicating that these proteins can facilitate biological interrogation. We subsequently selected the best IT setting for each method (MS2 500ms, RTS-MS3 750ms, RETICLE 750ms) and compared the quantified proteins (**Fig. 3E**). The results showed that most of the proteins were quantified across all methods and that RETICLE 750ms provided the highest number of additional proteins. To exemplify the individual advantages of the RTS methods, we investigated the quantification of the 40S and 60S ribosomal proteins, as these proteins should likely be co-expressed across single-cells. In the 1,076 commonly quantified proteins, 71 ribosomal protein were quantified and only RETICLE 750ms provided four additional ribosomal proteins from its 351 unique proteins. Comparing the pairwise correlations of these ribosomal proteins (**Fig. 3F**) showed that the expression profiles measured with RTS-MS3 750ms correlated the most and that both MS2-based methods performed similarly. Furthermore, the expression profiles from the additional proteins provided by RETICLE 750ms correlated well with the common proteins, supporting their accurate quantification. Taken together, these results confirm the observations from the diluted standard in the setting of a real scMS experiment and furthermore exemplify how the improved quantitative accuracy of RTS-MS3 or the increased proteome coverage of RETICLE is beneficial for scMS experiments.

**Figure 3.**
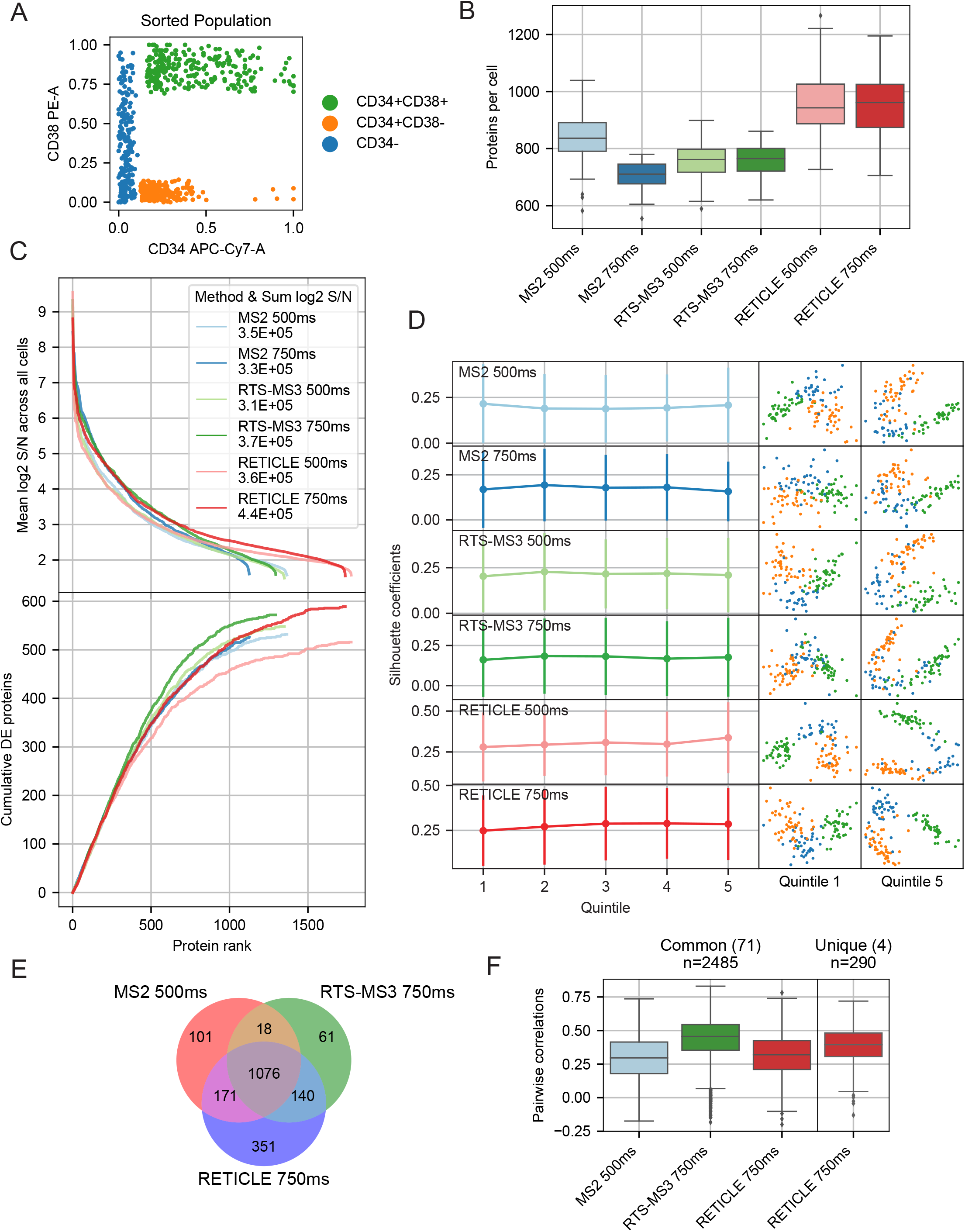
Comparison of acquisition methods using scMS samples. A) FACS strategy to sample single cells from three different populations of the OCI-AML8227 cell-culture system. B) Number of proteins quantified in each cell per method. C) Top: Distribution of S/N across proteins quantified in each method. For each method, proteins were ranked by their mean log2 S/N across all cells (different proteins for each method). Bottom: Cumulative distribution of DE proteins using the same protein ranking as top. DE analysis was performed on CD34+CD38+ population versus the rest of the cells. D) PCA analysis of the scMS dataset of each method with proteins from different rank bins. Proteins were ranked as in C and binned into quintiles. Missing values were imputed and protein expression was scaled. Separation of the different populations was measured by the silhouette coefficients of all cells in PCA space (first 10 PCs). Points show means and bars show standard deviation. PC1 & PC2 are shown on cell-scatterplots on the right. E) Venn diagram of proteins quantified in the three best methods. F) Pairwise correlations of expression profiles of ribosomal proteins (40S & 60S) across single cells in each method. Pearson correlation was calculated from imputed and scaled protein expression.

**Table 1.**
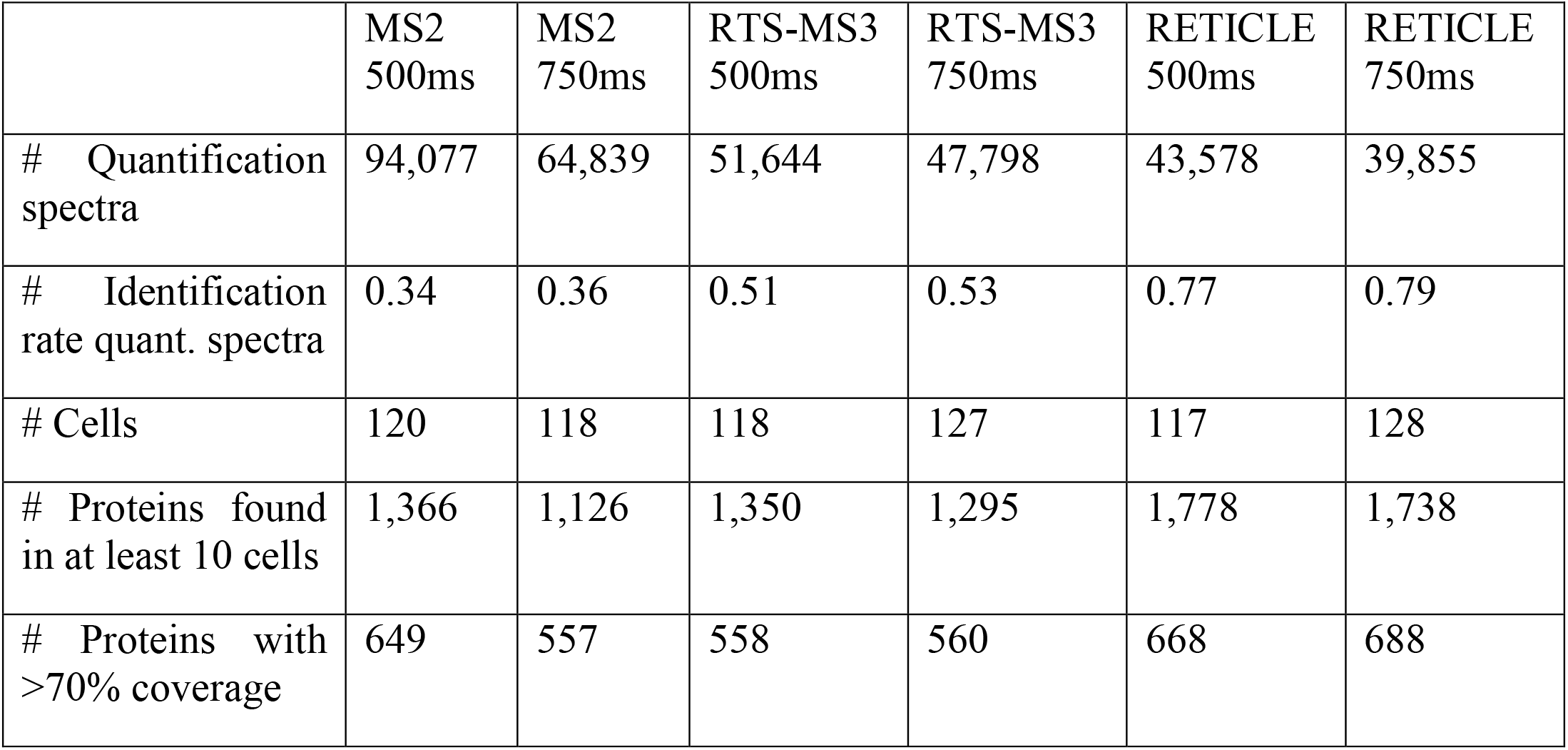
Comparison of scMS samples.

|  | MS2<br>500ms | MS2<br>750ms | RTS-MS3<br>500ms | RTS-MS3<br>750ms | RETICLE<br>500ms | RETICLE<br>750ms |
| --- | --- | --- | --- | --- | --- | --- |
| # Quantification spectra | 94,077 | 64,839 | 51,644 | 47,798 | 43,578 | 39,855 |
| # Identification rate quant. spectra | 0.34 | 0.36 | 0.51 | 0.53 | 0.77 | 0.79 |
| # Cells | 120 | 118 | 118 | 127 | 117 | 128 |
| # Proteins found in at least 10 cells | 1,366 | 1,126 | 1,350 | 1,295 | 1,778 | 1,738 |
| # Proteins with >70% coverage | 649 | 557 | 558 | 560 | 668 | 688 |

## DISCUSSION

Here we have shown that scMS data acquisition using the Tribrid design in combination with RTS can result in considerable improvements of scMS datasets and that the two RTS-based acquisition methods presented here have different advantages. RTS-MS3 provided a similar proteome coverage to MS2 at a much higher quantitative accuracy, whereas RETICLE resulted in higher proteome coverage. Thus, RETICLE could be especially useful for high throughput applications, such as building single-cell atlases, whereas RTS-MS3 could provide the accuracy needed to model protein networks^14^. To implement RTS-MS3 or RETICLE for scMS, we note that depending on the sample preparation, chromatographic setup and the level of boosting, ion injection times for the LIT and OT should be adapted to reach optimal performance. Furthermore, cycle times between FAIMS switching should be adapted to the chromatographic peak widths. Moreover, we note that in some cases, RETICLE can benefit from an intensity threshold filter to trigger LIT scans. Furthermore, as all methods showed similar performance at high protein S/N, it could be beneficial to regulate the IT using automatic gain control (AGC) to decrease acquisition time spent on highly abundant precursors. Finally, we hope our results inspire further pursuit of intelligent data acquisition strategies, with a lot of potential to be gained in the realm of single-cell proteome analysis.

## ABBREVIATIONS

AGC: Automatic gain control
AML: Acute myeloid leukemia
CID: Collision-induced dissociation
DE: Differential expression
HCD: Higher-energy C-trap dissociation
LC-MS: Liquid chromatography coupled mass spectrometry
LIT: Linear ion trap
LSC: Leukemic stem cells
OT: Orbitrap
PROG: Progenitor
RTS: Real-time search
S/N: Signal to noise ratio
scMS: Mass spectrometry-based single-cell proteomics
SPS: Synchronous precursor selection

## Data Availability

The mass spectrometry proteomics data will be deposited to the ProteomeXchange Consortium via the PRIDE^25^ partner repository. All data will also be available from the corresponding authors.

## Code availability

The complete analysis is available on Github (github.com/bfurtwa/RETICLE).

## Author Contributions

B.F. and E.M.S conceived the RETICLE method, designed and carried out experiments, and wrote the manuscript. B.F. conducted data analysis. N.Ü. performed FACS sorting and sample preparation. K.M, R.H, D.L.F and V.Z helped develop the instrument methods, and contributed to the overall functioning of the method. B.T.P. and E.M.S. oversaw the study. All authors proofread and contributed to the manuscript.

## Acknowledgments

This work was supported by the Independent Research Fund Denmark, the Svend Andersen Foundation, the Candys Foundation, the Danish Cancer Society and through a center grant from the Novo Nordisk Foundation (Novo Nordisk Foundation Centre for Stem Cell Biology, DanStem; Grant Number NNF17CC0027852). B. Furtwängler is the recipient of a fellowship from the Novo Nordisk Foundation as part of the Copenhagen Bioscience Ph.D. Programme, supported through grant NNF19SA0035442. We thank the BRIC FACS facility for support with the flow cytometry work. We thank the DTU Proteomics Facility for support with the MS analyses.

---

**Supplementary Figure 1.**
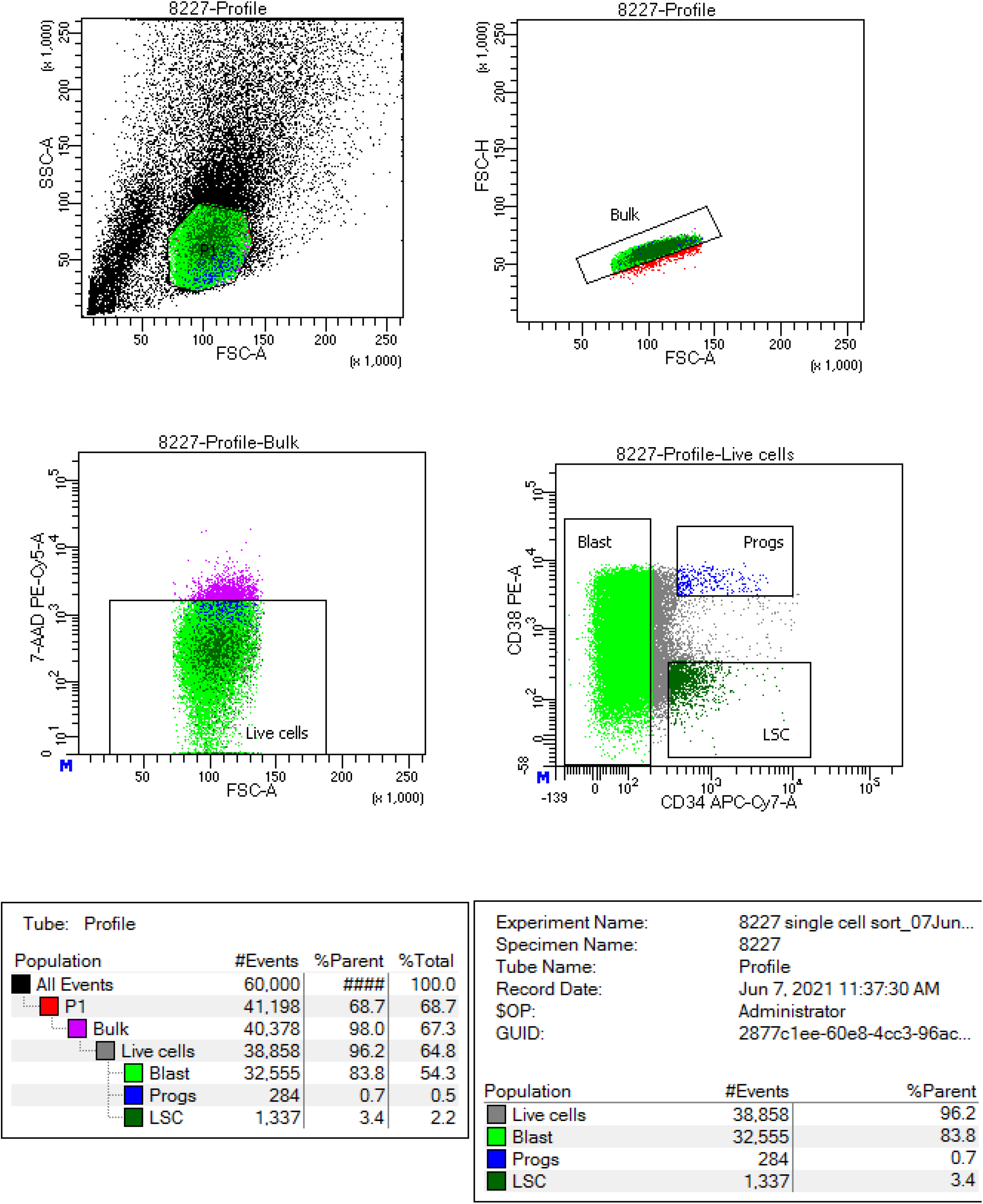
FACS strategy for single-cell sort of OCI-AML8227 cell-culture system.

**Supplementary Figure 2.**
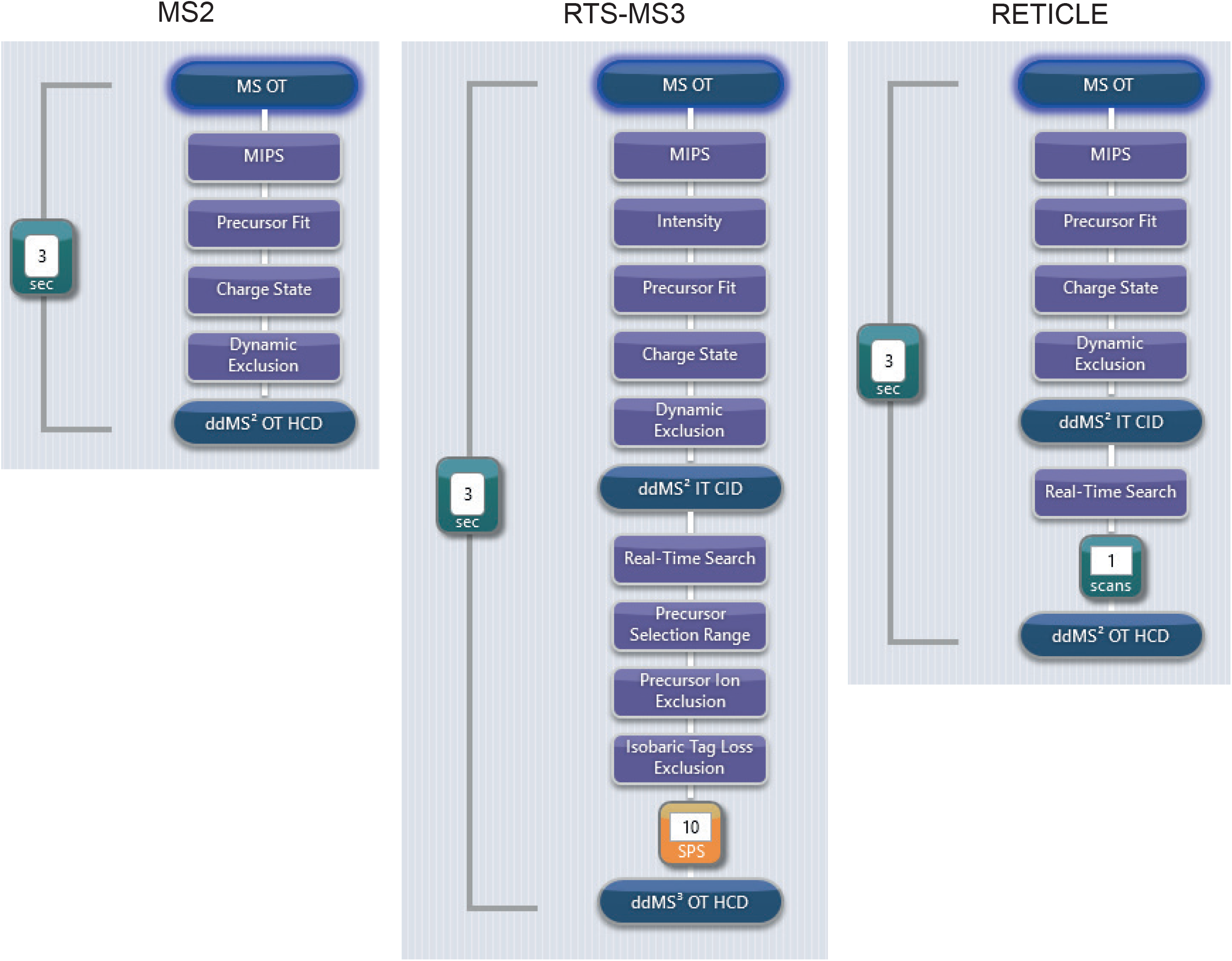
Method trees of the three acquisition methods.

**Supplementary Figure 3.**
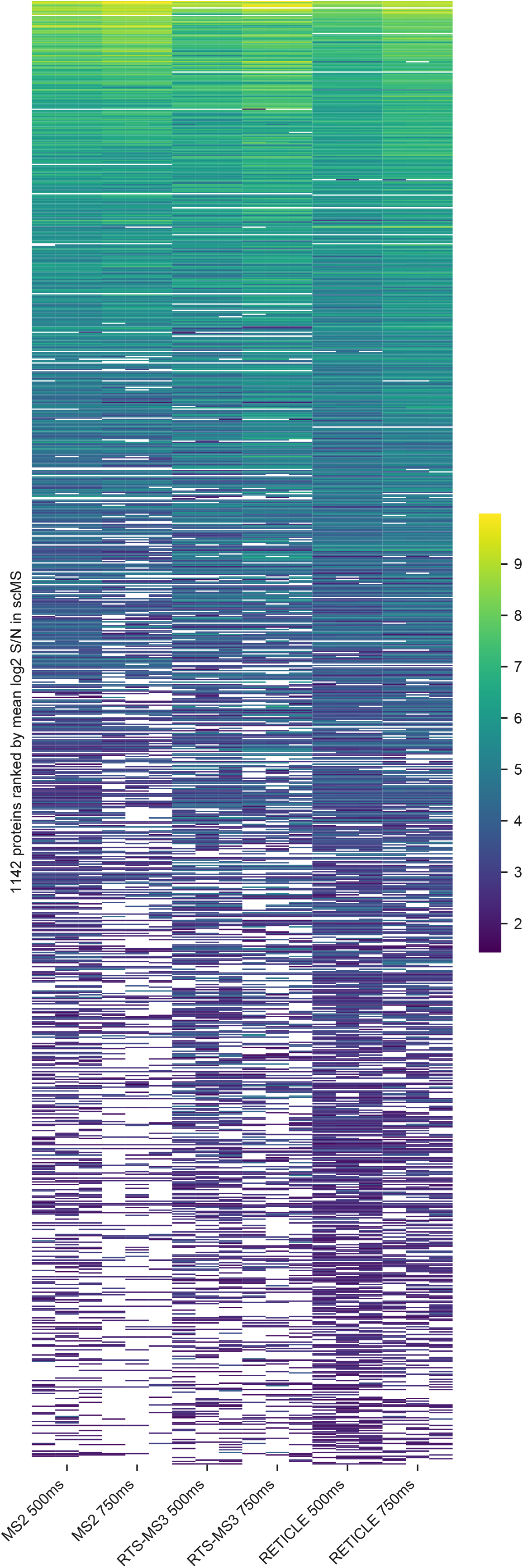
Full heatmap version of Figure 2A. Protein expression matrix of each method measured in triplicates. Proteins in the rows were sorted by the mean log2 S/N across all methods and each individual row shows the same protein with its mean log2 S/N across all cells in each LC-MS run.

